# Gaze Graphs in spatial navigation: visual behavior during free exploration explains individual differences in navigation performance

**DOI:** 10.1101/2025.10.28.684761

**Authors:** Jasmin L. Walter, Vincent Schmidt, Sabine U. König, Peter König

## Abstract

Vision is an important sense for spatial navigation, yet little is known about the role of visual behavior during spatial knowledge acquisition and its link to spatial navigation performance.

In this study, participants freely explored the immersive virtual reality city of Westbrook while eye-tracking data were collected. Subsequently, participants’ spatial knowledge was tested in an immersive pointing-to-building task. To address the freedom of movement during the spatial exploration, we transformed the eye-tracking data into gaze graphs and applied a graph-theoretical analysis. We assessed gaze graph architecture and identified gaze-graph-defined landmarks, finding that their properties were consistent with key results from our earlier work.

Furthermore, we investigated which factors contribute to the observed individual differences in spatial navigation performance. We found that global gaze graph measures, specifically gaze graph diameter, could explain 40% of the observed individual differences in participants’ mean task performance. In comparison, a model based on participants’ responses to a self-report questionnaire was not predictive of such differences. Consequently, we suggest that visual behavior, as reflected in global gaze graph measures, is a strong predictor of individual performance differences.

Our results not only contribute to a better understanding of landmarks and individual differences in spatial navigation but also outline the potential of vision and gaze-graph-based spatial navigation research.

## Introduction

The ability to orient and navigate within a new environment, for example, learning to find our way in a new city or building, is an essential skill in everyday life. As such, people typically navigate in noisy and complex environments, especially in cities and other urban spaces. However, despite spatial navigation having been studied for many decades, only more recently has the focus shifted to more naturalistic and complex settings (Bae et al., 2024; Delaux et al., 2021; Ishikawa et al., 2008; Ishikawa & Montello, 2006; Kiefer et al., 2014; Ohm et al., 2014, 2017). Especially the advancement in virtual reality (VR) technology over the last few years has offered new opportunities to combine complex and more naturalistic conditions with experimental control (Delaux et al., 2021; He et al., 2023; König et al., 2021; Schinazi et al., 2023; Varshney et al., 2024; Wunderlich & Gramann, 2021).

However, even with the rise in the use of more naturalistic and immersive experimental setups, reliably assessing and measuring active spatial learning and navigation remains challenging. For example, while more naturalistic setups might be utilized for participants’ spatial learning and exploration of an environment, they are often followed by tests that are conducted outside the initial environment, requiring a switch in modalities, e.g. (Ishikawa & Montello, 2006; König et al., 2021). Similarly, classical self-report questionnaires are often applied to identify differences in spatial strategy preferences and spatial navigation ability, but they are not always predictive of participants’ spatial navigation performance (Credé et al., 2019; König et al., 2019, 2021; Wunderlich & Gramann, 2021). In addition, while movement patterns during spatial navigation have recently started to receive more attention (Brunec et al., 2023; Sánchez Pacheco et al., 2025), our understanding of the participants’ continuous behavior during spatial learning, active orientation, and navigation remains severely limited. Overall, despite the methodological advances, current approaches remain limited in their ability to capture the continuous behavioral processes underlying spatial knowledge acquisition and linking them to navigation performance.

Furthermore, despite the importance of vision for spatial navigation (Chan et al., 2012; Ekstrom, 2015; Ekstrom & Isham, 2017; Foo et al., 2005) and common spatial concepts, we know only little about human visual behavior during spatial learning and navigation. For example, most landmark definitions are based on visible properties (Chan et al., 2012; Ekstrom, 2015; Ekstrom & Isham, 2017; Foo et al., 2005), yet we know only little about active visual behavior with respect to landmarks during both spatial learning and navigation. In general, landmarks play an important role in the formation of spatial knowledge (Presson & Montello, 1988; Siegel & White, 1975) and in spatial navigation (Chan et al., 2012; Foo et al., 2005; Richter & Winter, 2014; Steck, 2000; Yesiltepe et al., 2021). However, their exact function in the process of spatial knowledge acquisition and the formation of the mental map is still under discussion. Additionally, it remains unclear why certain buildings are used as landmarks and not others. Also, we know little about the role of environmental cues and city features beyond landmarks for spatial knowledge formation and differences in spatial navigation performance. Therefore, investigating visual behavior during spatial exploration with respect to environmental cues and landmarks could yield new insights into the role of landmarks and spatial learning in general.

As an initial step to address these challenges, we investigated free visual behavior during spatial learning in a complex immersive setting in our earlier work (Walter et al., 2022). Specifically, we collected eye-tracking data during active and free exploration of a virtual city using an immersive virtual reality setup. Since classical eye tracking analysis algorithms were not suitable to address participants’ freedom of movement during the free exploration of the city, we proposed a new analysis approach utilizing graph theory. Specifically, we transformed the eye tracking data into gaze graphs, allowing us to investigate general patterns in visual behavior across all participants while addressing the behavioral variability due to participants’ freedom of movement. The results of the graph-theoretical analysis offered new insights into vision during spatial knowledge acquisition. Importantly, we identified a subgroup of buildings in the city that were consistently important in participants’ visual network during the free exploration of the city and were subsequently classified as gaze-graph-defined landmarks. In addition, we also discovered that these gaze-graph-defined landmarks were preferentially connected to each other, suggesting that they might serve as a network of reference points, thus offering new insights into survey knowledge formation. However, since this was the first study of its kind, the question remains how stable the gaze graphs and gaze-graph-defined landmarks are in their characteristics and properties. Moreover, without connecting the observed visual patterns with the accuracy of participants’ survey knowledge and allocentric representation, the role of visual behavior in spatial knowledge formation cannot be fully addressed and investigated. Therefore, a new experimental approach was needed to directly link visual behavior during spatial learning to spatial navigation performance.

When investigating spatial navigation performance, a consistent finding is the considerable presence of large individual differences in navigation performance. In general, individual differences in spatial navigation are ubiquitous in the existing literature and can be observed in a variety of tasks (Boone et al., 2019; Ekstrom & Isham, 2017; Gramann, 2013; He et al., 2023; Hegarty et al., 2018, 2023, 2023; Ishikawa, 2023; Ishikawa & Montello, 2006; Maxim & Brown, 2023; Newcombe, 2018; Newcombe et al., 2023; Schinazi et al., 2023; Spiers et al., 2023; Weisberg et al., 2014; Weisberg & Newcombe, 2014, 2016, 2018; Zanchi et al., 2022). They are especially prevalent in people’s ability to learn a new environment and form a map-like knowledge (also referred to as survey knowledge or allocentric representation) (Hegarty et al., 2018; Ishikawa, 2023; Ishikawa & Montello, 2006; Newcombe et al., 2023; Weisberg et al., 2014; Weisberg & Newcombe, 2016, 2018). In general, a multitude of factors have been found to contribute to the observed individual differences. Specifically, there is evidence that suggests that people possess differing levels of spatial (navigation) capabilities, indicated by matching performance differences in other spatial reasoning tasks (Hegarty et al., 2018) and the size of brain regions (Maguire et al., 2000). Another factor often considered is gender (Boone et al., 2019; Brunyé et al., 2018; Coutrot et al., 2018; Lawton, 1996; Nazareth et al., 2019; Schinazi et al., 2023; Spiers et al., 2023). In addition, environmental factors like the spatial entropy of the city participants grew up in (Coutrot et al., 2022) or cultural and language differences (Coutrot et al., 2018; Haun et al., 2011; Hund et al., 2012; Spiers et al., 2023) have been found to contribute. Furthermore, some of the observed differences can be manipulated and reduced with specific instructions (Boone et al., 2019) or motivation introduced by a monetary penalty (Schinazi et al., 2023). Overall, individual differences appear to be rooted in multiple developmental and environmental factors (Gramann, 2013; Newcombe et al., 2023). Altogether, this suggests that visual behavior could also contribute to and reflect the observed individual differences. Therefore, exploring the link between visual behavior during spatial learning and individual differences in spatial navigation performance could offer a new perspective into this phenomenon and the role of visual behavior for spatial navigation in general.

In the following study, we combine a long free exploration period of the new virtual city of Westbrook with immersive pointing tasks conducted in the same environment using an immersive virtual reality setup with eye tracking functionality (Schmidt, König, Dilawar, Sánchez Pacheco, et al., 2023). The experimental design not only enables an investigation of the visual behavior during spatial exploration and learning, but also allows us to link participants’ exploration behavior with their spatial navigation performance within the same urban environment, consequently extending and improving the experimental design from our earlier work (Walter et al., 2022). The current study is part of a larger experiment framework and focuses on the analysis of the visual behavior and linking it to participants’ task performance. An analysis of the behavioral and qualitative self-reported changes related to spatial learning and a comparison to a participant group trained in an augmented condition is published in (Schmidt, König, Dilawar, Sánchez Pacheco, et al., 2023). Here, we transform the collected eye-tracking data during the free exploration of the environment into gaze graphs, apply the same graph-theoretical analysis approach from our earlier work (Walter et al., 2022), and link general patterns in visual attention during spatial exploration to participants’ pointing task performance. Specifically, we investigate the general architecture of gaze graphs, identify gaze-graph-defined landmarks, assess their characteristics, and ultimately discuss the consistency of the graph-theoretical results in both studies. Furthermore, we link participants’ visual behavior during spatial exploration and their self-reported navigation abilities to spatial knowledge accuracy by modeling task performance differences with FRS questionnaire responses and global gaze graph measures. Overall, we investigate the characteristics and consistency of visual behavior during spatial knowledge formation and the link to spatial navigation performance, specifically survey knowledge and the mental map.

## Results

In the study, 26 participants used an immersive virtual reality (VR) setup (Figure 1a) to freely explore the virtual city of Westbrook (Figure 1b–c), and subsequently completed an immersive pointing-to-building task within the same environment (Schmidt, König, Dilawar, Sánchez Pacheco, et al., 2023). All participants wore the head-mounted virtual reality headset Vive Pro Eye. Eye-tracking data was collected with a mean sampling rate of 90Hz. In the first part of the study, participants explored the city for five 30-minute recording sessions, thus for a total of 2.5 hours. The participants did not receive any other task than to explore the city and take a picture of all buildings that had street art painted on them using their controller buttons. Every 10 minutes, the eye tracker was validated and, if necessary, recalibrated. All participants were seated on a swivel chair that allowed them to fully rotate their bodies while they controlled their walking speed with the controller (Figure 1a). After the exploration phase, the participants performed the pointing-to-building task, in which they were teleported to different starting locations in the city and asked to point to a specific building with their controller. In addition, at the beginning of the experiment, participants were asked to complete the FRS questionnaire on spatial strategies, containing self-report questions on participants’ use of spatial navigation strategies in everyday situations and perceived orientation ability (Münzer & Hölscher, 2011).

**Figure 1:**
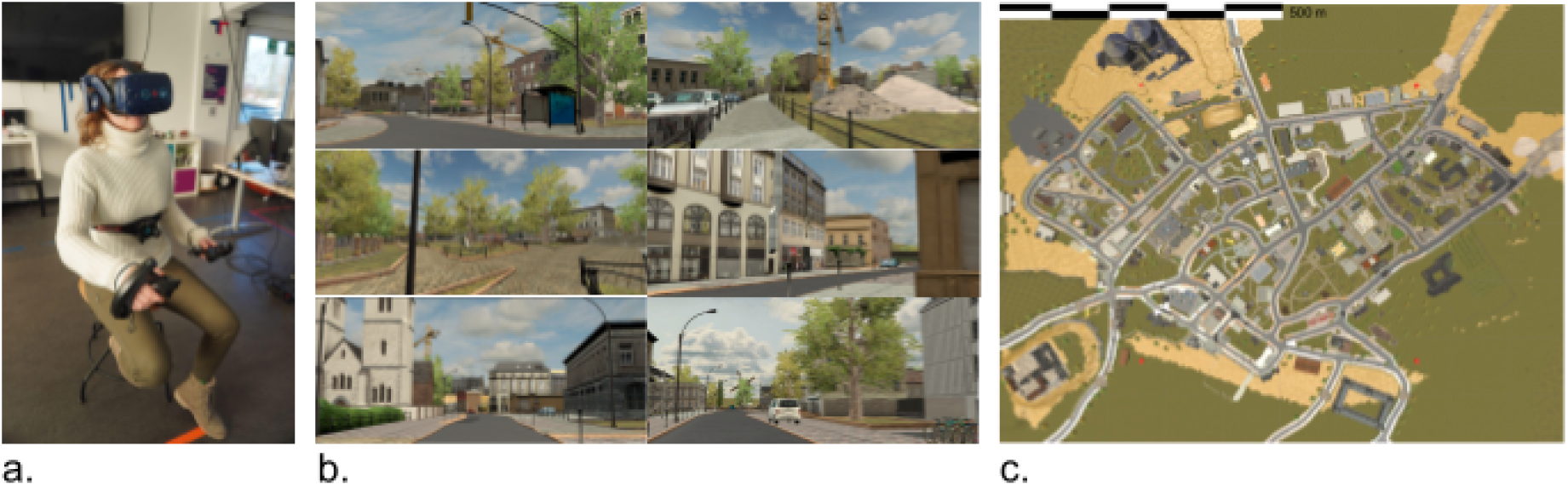
Experimental Setup and the virtual city of Westbrook. **a)** Participant in the immersive virtual reality setup. **b)** Impressions from a first-person perspective when exploring the virtual city of Westbrook. **c)** The map of Westbrook, a fictional town inspired by the city of Baulmes in Switzerland.

### Gaze graph creation

To investigate visual behavior during 2.5 hours of free spatial exploration in a large urban environment, the eye tracking analysis needed to address the variability of the participants’ behavior resulting from their freedom of movement. Specifically, participants were able to move their heads and eyes freely without any restrictions, in addition to freely walking within the whole city (approximately 996m x 758m ∼ 75.5 hectares) while only being limited to the road network and other walkable areas (other narrow stone or grass paths, parking areas, grass patches, etc.). Consequently, participants were walking in different areas of the city and looking at different objects in the environment at any time during the experiment, as exemplified by the walking paths (Figure 2a) and gaze locations (Figure 2b) of six example participants within the first five minutes of the experiment. Therefore, an abstraction process was needed to identify common patterns of visual behavior without the behavioral variation introduced by different walking decisions. We therefore applied the graph-theoretical analysis approach from our earlier work (Walter et al., 2022) and transformed the eye-tracking data during the exploration phase of the experiments into gaze graphs. By creating gaze graphs, we reduce the dimensionality of the eye-tracking data, thereby making the relevant features of the data accessible and excluding variation due to participants’ freedom of movement. We suggest that transforming the eye-tracking data into gaze graphs appropriately addresses the freedom of participants’ behavior and allows us to investigate the underlying visual attention patterns during spatial exploration across participants.

**Figure 2:**
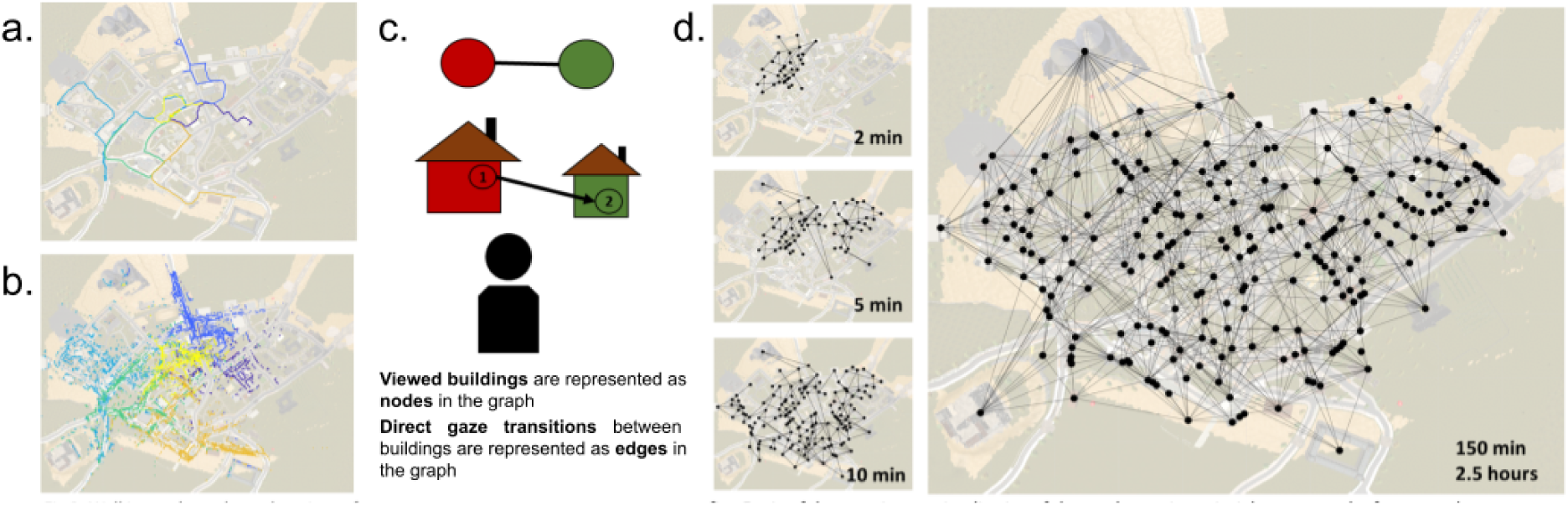
Gaze graphs creation. **a), b)** Walking paths (a) and gaze locations (b) of 6 example participants during the first 5 min of the experiment. Each color corresponds to one participant. **c)** Visualization of the gaze graph creation principle. **d)** Gaze graph of an example participant after 2 min, 5 min, 10 min, and the full 150 min / 2.5 hours of free exploration.

We created gaze graphs from the eye-tracking data in a series of standardized steps. First, a cleaning and pre-preprocessing pipeline was applied (see Data pre-processing and graph-theoretical analysis). Then, the recorded eye-tracking data was transformed into gaze graphs using the following process. Each viewed building in the city was represented by a node in the gaze graph. Edges between nodes were created when participants made a direct gaze transition from one building to the other. For example, if a participant gazed at building 1 and directly continued with a gaze at building 2, this behavioral pattern resulted in two nodes representing Building 1 and Building 2, which were connected with an edge, representing the direct gaze transition between both buildings (Figure 2c). Importantly, the gaze graph was solely based on the eye tracking information. Thus, the physical location of the participants in the city was not represented in the gaze graph. Furthermore, the gaze graph was undirected and unweighted. Consequently, the gaze graph only contained information about whether a participant performed a direct gaze transition between two buildings without maintaining any information about the directionality or the frequency of this gaze transition. In short, the 2.5 hours of eye-tracking data recorded in the exploration phase were reduced to one gaze graph for each participant (Figure 2d).

To identify general gaze graph characteristics and investigate common visual patterns between participants’ visual exploration of the city, we applied the graph-theoretical analysis pipeline from our earlier work (Walter et al., 2022). Specifically, we investigated whether the virtual city could be treated as one coherent city in the analysis and identified potential gaze-graph-defined landmarks and their connectivity. Since the present study and our earlier work (Walter et al., 2022) share both the graph-theoretical analysis of gaze graphs and a similar experimental design (e.g., free exploration in a virtual city), we list a detailed overview of both experiments and results in Table 3. In the discussion, the results of the graph-theoretical analysis in both experiments are compared, and similarities and differences are addressed. The following section will focus on the results of the graph-theoretical analysis based on the eye-tracking data and gaze graphs of this study.

### Gaze graph properties and spectral graph analysis

As a first step, we investigated whether the city of Westbrook was perceived as one coherent single city or as a set of loosely connected areas that need to be treated separately. In the following analysis, we applied a spectral graph analysis to investigate the gaze graph’s connectivity based on the eigenvalue spectrum of the Laplacian matrix. To calculate the Laplacian matrix, we subtracted the adjacency matrix from the degree matrix of the gaze graph. We then assessed the second smallest eigenvalue of the Laplacian matrix, the Fiedler value, as a measure of the overall connectivity of the graph, and found a mean Fiedler value of 0.353 (*SD* = 0.093) over all graphs.

As a second step, we identified the Fiedler vector, which is defined as the eigenvector corresponding to the second-smallest eigenvalue. The Fiedler vector assigns one value to each node that can be used to partition the graph into two clusters based on whether the node value of the vector is positive or negative. Furthermore, the Fiedler vector gives insights into the interconnectivity of the nodes. Specifically, the further the Fiedler vector entries are away from 0, the higher the connectivity of the particular node within its cluster. When assessing the Fiedler vector of all gaze graphs, we found many Fiedler vector values around 0, indicating that a considerable number of nodes do not belong to either cluster. As a consequence, the gaze graphs did not appear to separate clearly into two natural clusters. Furthermore, by sorting the adjacency matrix according to the sorting index of the Fiedler vector, the potential clusters of the graph can be visualized. In this spectral matrix visualization, a graph with easy-to-separate clusters would display a modular block pattern. However, we were not able to find this block pattern in any of the gaze graph visualizations. Consequently, gaze graphs lacked a clear modular pattern, thus, we found no clear separation between clusters.

Finally, we partitioned the gaze graphs based on the sparsest cut approximated by the Fiedler vector and assessed the connectivity and composition of the graphs before and after the spectral partitioning. Before the partitioning, the uncut gaze graphs contained, on average, 1295.1 edges (*SD* = 206.3) with a density of 0.046. Consequently, on average, 4.6% of all possible edges were instantiated. Partitioning the gaze graph into two components using the sparsest cut approximated by the Fiedler vector (i.e., separating the graph based on the positive and negative values of the Fiedler vector) required cutting 122.8 edges on average (*SD* = 29.5), resulting in a cut of 9.39% of the graph’s edges on average (*SD* = 1.2%). The resulting two clusters had a similar average number of edges (edges cluster 1: *M* = 584.7, *SD* = 155.7; edges cluster 2: *M* = 587.7, *SD* = 212.0). After the partitioning, the mean intra-cluster density amounted to 0.0873 (*SD* = 0.0133) and the inter-cluster density averaged 0.0092 (*SD* = 0.0024). These numbers indicated an increase in intra-cluster density with a simultaneous decrease in inter-cluster density after the partitioning. Nevertheless, when interpreting these values, it was important to consider that in order to separate the graph into clusters, a fair portion of edges had to be cut. Taken altogether, the results of the full spectral partitioning analysis indicated that the gaze graphs had no clear modular structure, and we found no clear separation between clusters. Thus, the graph could not be partitioned without cutting a considerable number of edges. Consequently, the city of Westbrook was visually perceived as a unit rather than a set of separate city blocks, indicating that the virtual city could be treated as one coherent environment in the following analysis.

### Gaze-graph-defined landmarks

To identify possible gaze-graph-defined landmarks, we assessed the distribution of node degree centrality for all participants (Figure 3a). Our results showed moderate variation of the node degree averaged across buildings per participant (*M* = 10.62, *SD* = 1.69). The correlation of buildings’ node degrees between participants resulted in a mean coefficient of 0.74 (*SD* = 0.12) (Figure 3c). Thus, the viewing behavior of different participants as captured by the buildings’ node degrees appeared to be comparable across participants.

**Figure 3:**
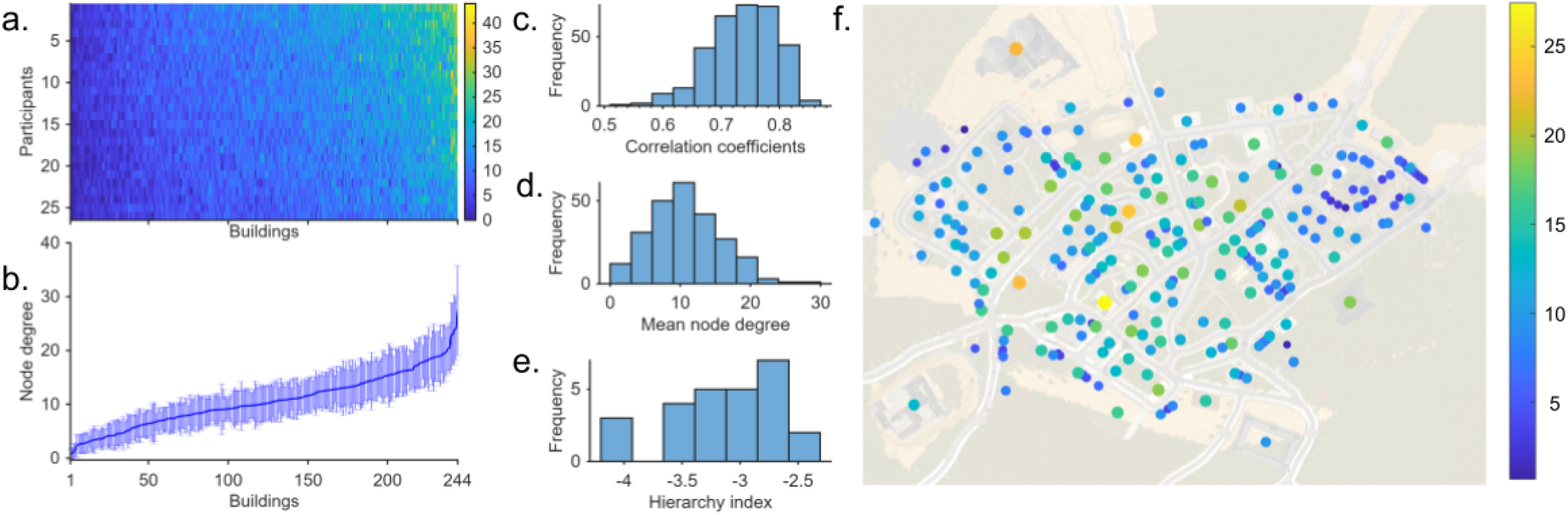
Node degree centrality. **a)** all node degree values of all participants (vertical axis) and all buildings (horizontal axis). All values are sorted with increasing mean node degree from bottom to top along the vertical axis and from left to right along the horizontal axis. **b)** The mean node degree of each building averaged over participants matching the sorting order in a). **c)** Histogram of the pairwise correlation coefficients. **d)** Histogram of the mean node degree values of each building, averaged over participants**. e)** Histogram of the hierarchy index of all gaze graphs. **f)** The locations of all buildings on top of the map are color-coded with the buildings’ mean node degree.

As a next step, we compared the node degrees of different buildings. When averaged across participants, our results showed a large variation in the buildings’ node degrees (*M* = 10.62, *SD* = 4.96, Figure 3b). In addition, the hierarchical configuration of the gaze graphs, as characterized by the hierarchy index (*M* = −3.1, *SD* = 0.47) (Figure 3e), resulted in a small number of nodes with very high degree values for each participant, thereby highlighting the importance of these highly connected nodes within the gaze network. Further examination of the distribution of the buildings’ mean node degrees revealed a small subset of buildings that clearly stood out (Figure 3d), suggesting that important nodes were consistent across participants. Next, we applied the node degree threshold for gaze-graph-defined landmarks, defined as two standard deviations over the mean node degree averaged over all participants and buildings (Walter et al., 2022). Five buildings exceeded this threshold (mean + 2 SD = 20.53). Therefore, considering the hierarchical configuration of the gaze graphs and applying the node degree threshold for gaze-graph-defined landmarks, we identified five gaze-graph-defined landmarks.

### Location and appearance of gaze-graph-defined landmarks

To further characterize the gaze-graph-defined landmarks, we examined their locations and appearances in the environment. We found that all five gaze-graph-defined landmarks had a very distinct visual appearance and were always located at a crossing, therefore making them visible from multiple angles (Figure 4). In addition, the five gaze-graph-defined landmarks were distributed across the central-western area of the city. Therefore, all identified gaze-graph-defined landmarks matched the appearance and location characteristics of the gaze-graph-defined landmark definition (Walter et al., 2022).

**Figure 4:**
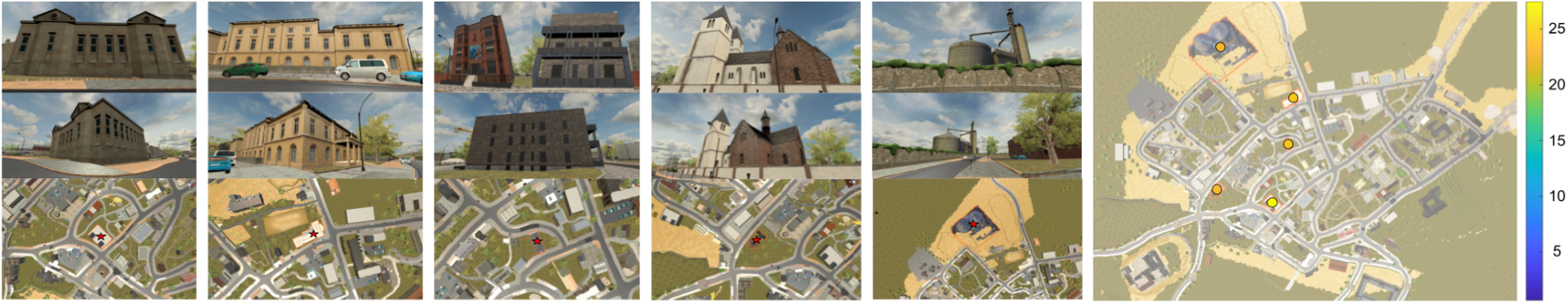
Gaze-graph-defined landmarks: Images of all gaze-graph-defined landmarks from multiple perspectives, with their location marked on the zoomed-in map and the whole city map with the marked gaze-graph-defined landmarks.

To assess the potential of gaze-graph-defined landmarks to be utilized for triangulation, directional cues, or beacons, we calculated their visibility with respect to the walkable area and weighted by the time participants spent in different parts of the city. First, we calculated the visibility of each building by identifying the location from which participants gazed at the buildings using a spatial resolution of 4 x 4 m. We included only locations in the city that were visited by at least one participant, although the majority of the walkable area was explored by all participants (Figure 5a). Consequently, a building was considered visible if at least one participant viewed the building from the respective location. In general, gaze-graph-defined landmarks were visible from 7.21% of the traversed area on average (*SD* = 0.57%) while other buildings were visible from 2.77% of the traversed area on average (*SD* = 1.47%). Therefore, gaze-graph-defined landmarks were visible from a much larger proportion of the city compared to other buildings. Furthermore, we found that in 29% of the traversed area, at least one gaze-graph-defined landmark was visible (Figure 5b). More specifically, in 7% of the traversed area, two or more gaze-graph-defined landmarks were visible at the same time. This could have allowed participants to triangulate their position using two or more gaze-graph-defined landmark cues. Correspondingly, in 22% of the traversed area, exactly one gaze-graph-defined landmark was visible, thus offering the basis for directional cues and beacon navigation (Figure 5b). In the remaining 72% of the city, no gaze-graph-defined landmark was visible (Figure 5b). Despite covering only about 29% of the traversed area, locations where at least one gaze-graph-defined landmark was visible accounted for 46.9% of exploration time. Of this time, 31.8% was spent where exactly one gaze-graph-defined landmark was visible, and 15.1% where two or more were visible. The remaining 53% was spent where no gaze-graph-defined landmarks were visible. Overall, gaze-graph-defined landmarks displayed an increased visibility compared to other buildings and offered the opportunity for several typical landmark uses like triangulation, directional cues, and beacon navigation across nearly one third of the walkable area, in which participants also spent nearly half of the exploration time.

**Figure 5:**
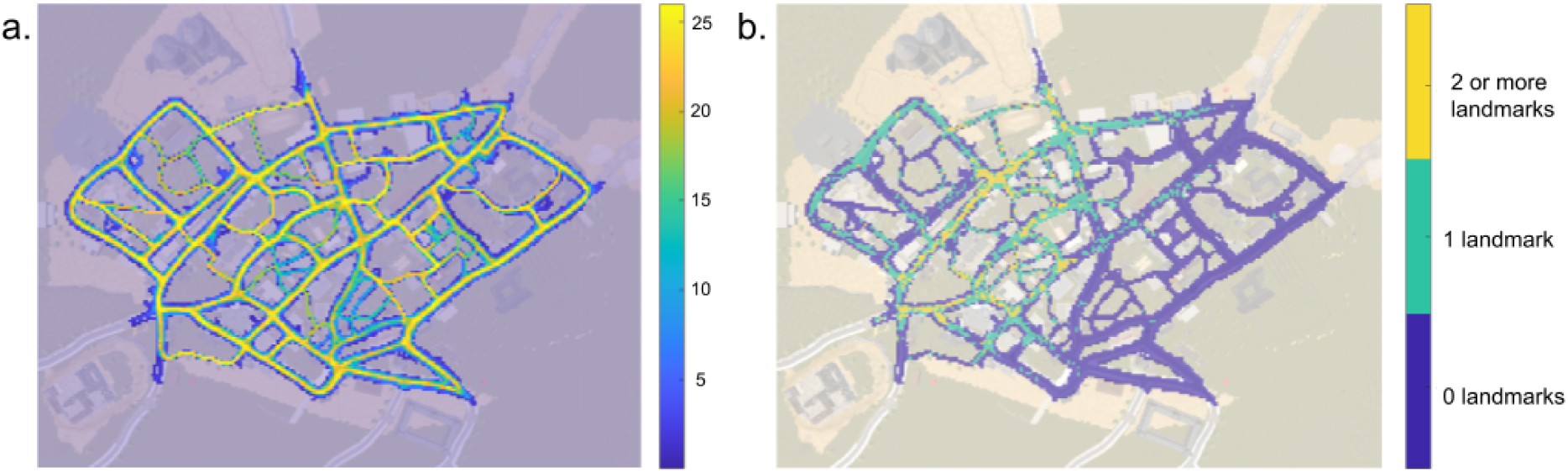
Walking paths and visibility of gaze-graph-defined landmarks. **a)** Walking paths of all participants, color coded by the number of participants visiting a specific area of the city (calculated with a 4×4m spatial resolution). **b)** Walking paths of the city, color coded from which areas 0, 1, 2 or more gaze-graph-defined landmarks were viewed by participants (4×4m spatial resolution).

### Interconnectedness of gaze-graph-defined landmarks

To identify whether the gaze-graph-defined landmarks are also preferentially connected to each other, we applied the rich club analysis pipeline (Walter et al., 2022). As a first step, we calculated the mean adjusted rich club coefficient to identify the connectivity of the whole gaze graph. The adjusted rich club coefficient compares the connectivity of a gaze graph with a random graph with matched node degree statistics. Therefore, we generated 1000 random graphs with the same number of nodes as the gaze graph and identified the 10 graphs that were most similar to the node degree distribution of the original graph using the two-sample Kolmogorov-Smirnov test. Next, we calculated the adjusted rich club coefficient by taking the ratio of the rich club coefficient of the gaze graph and the rich club coefficient for one of the random graphs, and repeated the process for all the remaining nine random graphs. By averaging all 10 adjusted rich club coefficients, we calculated the mean adjusted rich club coefficient, where exceeding a value of 1.0 indicates preferential connectivity between the nodes of the graph.

As a second step, we systematically removed low-degree nodes from the set of nodes considered for the mean adjusted rich club coefficient calculation. We found that with an increasing minimum node degree, the mean adjusted rich club coefficient showed a consistent increase across all participants (Figure 6a). That is, nodes with a higher node degree centrality were more densely interconnected than expected by chance. However, since the increasing node-degree threshold also resulted in fewer nodes being included in the calculation, the estimate’s uncertainty increased considerably with the increasing node-degree threshold. Even though the rich club coefficient does not have a clear cutoff point, we applied the threshold of one-sigma distance above the overall mean node degree as the highest value for reliably calculating the rich club coefficient (Walter et al., 2022). This resulted in the observation that the nodes with a node degree of 15 and higher were interconnected the most in the gaze graph compared to the corresponding random graphs. Consequently, buildings with a high node degree formed a rich club within the gaze graph.

**Figure 6:**
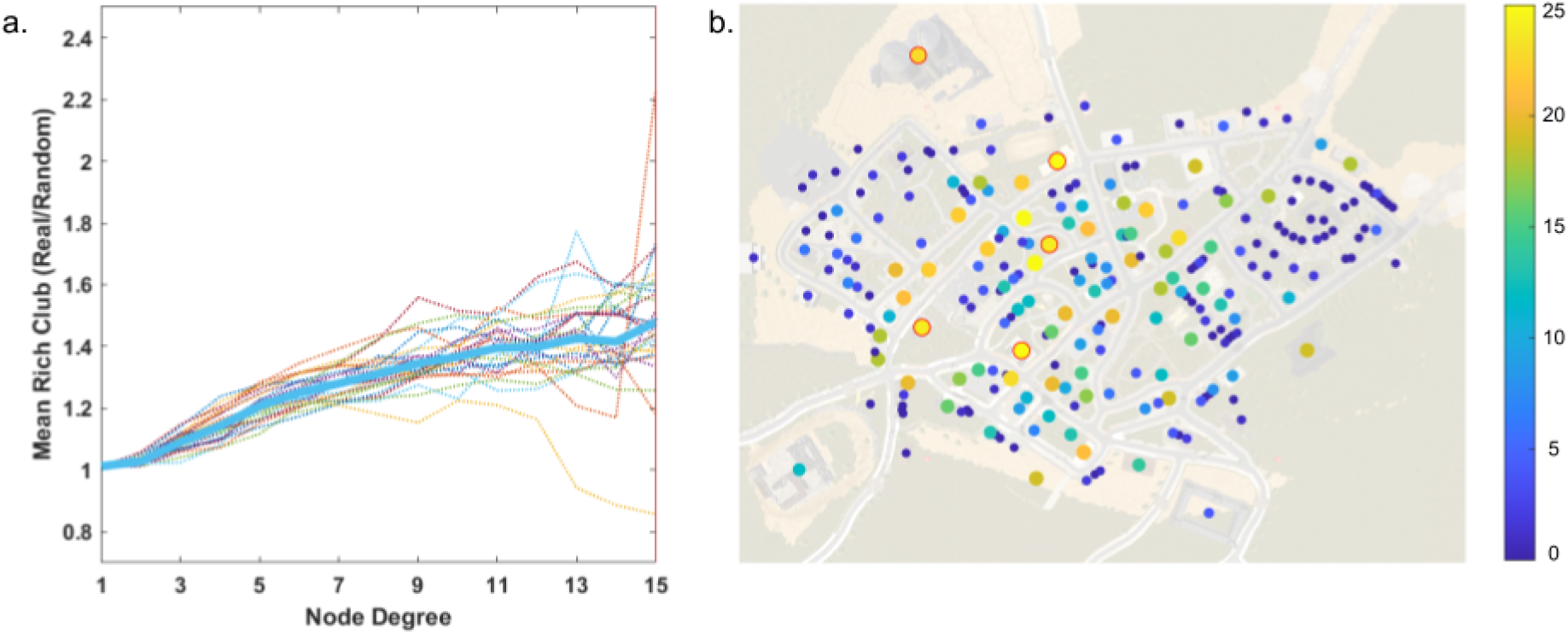
Rich club analysis. **a)** Mean adjusted rich club coefficient (y-axis) with increasing node degree included in the calculation (x-axis). The blue line marks the population mean of the individual mean adjusted rich club coefficient. **b)** All buildings are plotted on the map of Westbrook with the color and size reflecting how frequently the building appeared in the final rich club of individual participants (including nodes of 15 node degree centrality and higher). The five gaze-graph-defined landmarks are additionally marked with a red edge color.

As a final step in the pipeline, we identified which buildings appeared most often in the rich club across participants. Specifically, we counted the number of times buildings were part of the final set of nodes considered for calculating the adjusted rich club coefficient before the cut-off threshold, i.e., buildings with a node degree of 15 and higher, across participants. Then, we identified the buildings exceeding the threshold of two standard deviations above the mean count. These 14 buildings appeared most often in the strongest rich club across participants and included all five gaze-graph-defined landmarks (Figure 6b). Therefore, high node degree buildings and gaze-graph-defined landmarks were preferentially connected and part of the strongest rich club within the gaze graphs.

### Spatial navigation performance

To assess participants’ spatial knowledge, they performed an immersive pointing-to-building task after completing the free exploration of the city of Westbrook (Schmidt, König, Dilawar, Sánchez Pacheco, et al., 2023). In the pointing-to-building task, participants were asked to indicate the direction to specific buildings within the city. The task utilized eight task buildings and eight starting locations that were directly located in front of each task building (Figure 7b-c). The buildings were distributed evenly over the city (Figure 7b) and selected so that they were not visible from any other starting location. Participants were teleported to each of the starting locations in randomized order and rotated to a randomized degree during the teleportation. At each starting location, they were shown a picture of the target building and asked to point to the building using their controller. In the participant’s view, this was displayed as a green ray indicating the pointing direction (Figure 7a). The participants logged their pointing response with a button press. At each task location, the participants pointed to the remaining seven other buildings, resulting in 56 unique start and target building combinations. After completing the task, the participants repeated all trials in a newly randomized order. Overall, each participant performed 112 trials in the pointing-to-building task.

**Figure 7:**
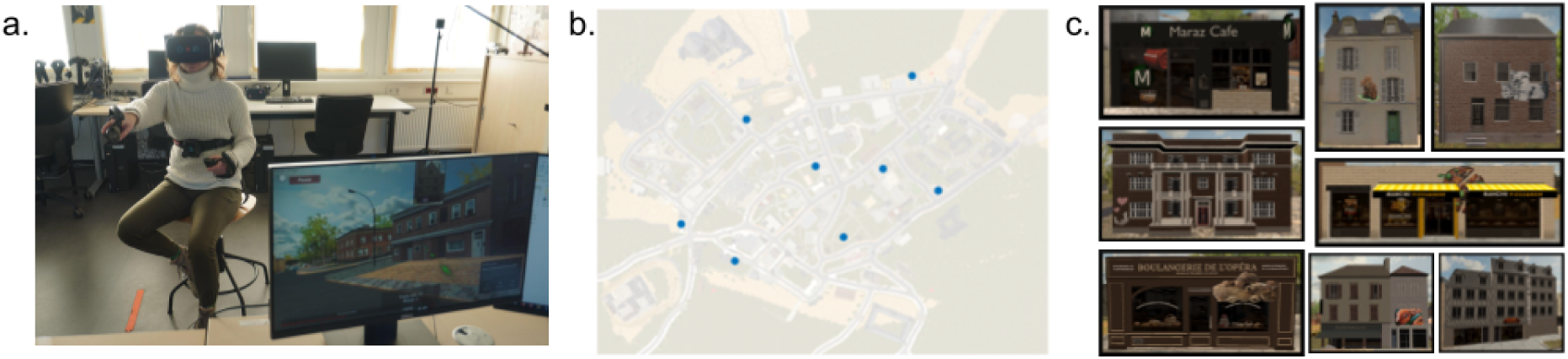
Pointing-to-building task. **a)** Experimental setup during the immersive pointing-to-building task. The VR view is visible on the computer screen, including the green pointing marker. **b)** Location of all eight task buildings within the city. **c)** Pictures of all eight task buildings.

We assessed participants’ performance by calculating the absolute pointing error deviation from the correct direction. Here, a pointing error of 0° represents the best possible task performance; a pointing error of 180° represents the worst possible task performance. In Figure 8a, all pointing errors for each participant and each unique start-target-building combination are displayed. The mean performance of participants varied considerably between the best-performing participant (pointing error: *M* = 15.2°) and the worst-performing participant (pointing error: *M* = 74.0°), resulting in a difference in mean pointing error of 58.8° between the best and worst-performing participants (Figure 8b). Thus, we found considerable variance in task performance between participants.

**Figure 8:**
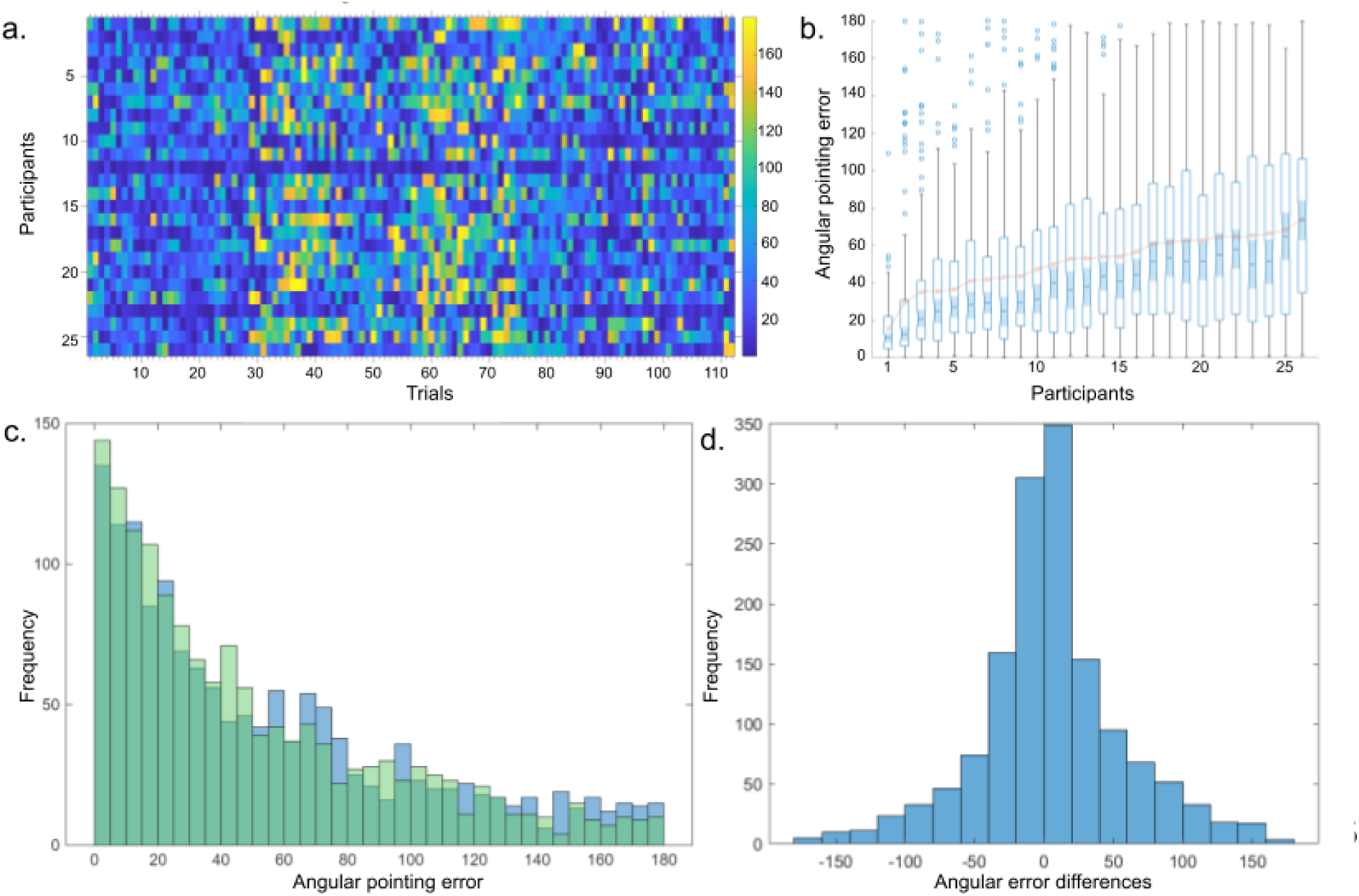
Task performance. **a)** All angular pointing errors of all 26 participants (vertical axis) and all 112 pointing trials (horizontal axis). **b)** Boxplot chart of participants’ angular pointing error, with the mean performance marked (orange overlay). **c)** Histogram of all pointing errors separated by trial order: blue = initial trials, green = repeated trials. **d)** Histogram of the deviation in pointing angle between the first and repeated trials.

Next, we assessed systematic differences in participants’ performance during the repetition trials, specifically, the performance differences between the initial and repeated presentations of each trial. When visually inspecting the pointing error distributions of the initial and repeated trials, they appeared to be quite similar (Figure 8c). Additionally, the distribution of the differences in pointing direction, for example, the change in the pointing direction between the initial and repeated response for the same trial, appeared to be normally distributed (Figure 8d). However, when performing a repeated-measures ANOVA to examine the effect of trial repetition order (initial answer and repeated answer) and trial identity (unique-start-target-building combination), we found a significant main effect of repetition order, *F*(1, 25) = 6.97, *p* = .014. In addition, we found a highly significant main effect of trial identity, *F*(55, 1375) = 7.11, *p* = 1.23 × 10⁻⁴⁴. The interaction between repetition order and trial identity was not significant, *F*(55, 1375) = 1.16, *p* = .201. On average, participants’ pointing errors decreased by 5.1° (*SD* = 9.9°) when repeating the trial a second time during the task repetition. To summarize, participants performed significantly better during the task repetition, showing an average 5.1° improvement in pointing accuracy.

### Modeling individual differences in spatial task performance

In the following, we investigated the relationship between individual differences in task performance and participants’ visual behavior during the free spatial exploration of the city and their self-assessment of their spatial navigation skills (FRS questionnaire (Münzer & Hölscher, 2011)). Since differences in pointing performance between initial and repeated trials cannot be explained by measures of spatial exploration (e.g., spatial learning or navigation ability), we first quantified how much of the total variance in pointing error could be attributed to trial repetition. The fraction of the variance that could not be explained by performance differences during the repetition trials served as a baseline for the maximum explainable variance under an ideal model.

To assess the fraction of variance attributed to performance differences between initial trials and their repetitions, we first calculated the intra-participant variance for each unique start-target building combination. We found that intra-participant variance of each unique start-target building combination accounted for 65.64% of the total variance across all pointing trials (Figure 9a). As a consequence, when modeling participants’ performance across trials, even a best-performing model using variables related to participants’ spatial exploration and general spatial abilities could only explain at most 34.36% of the total variance.

**Figure 9:**
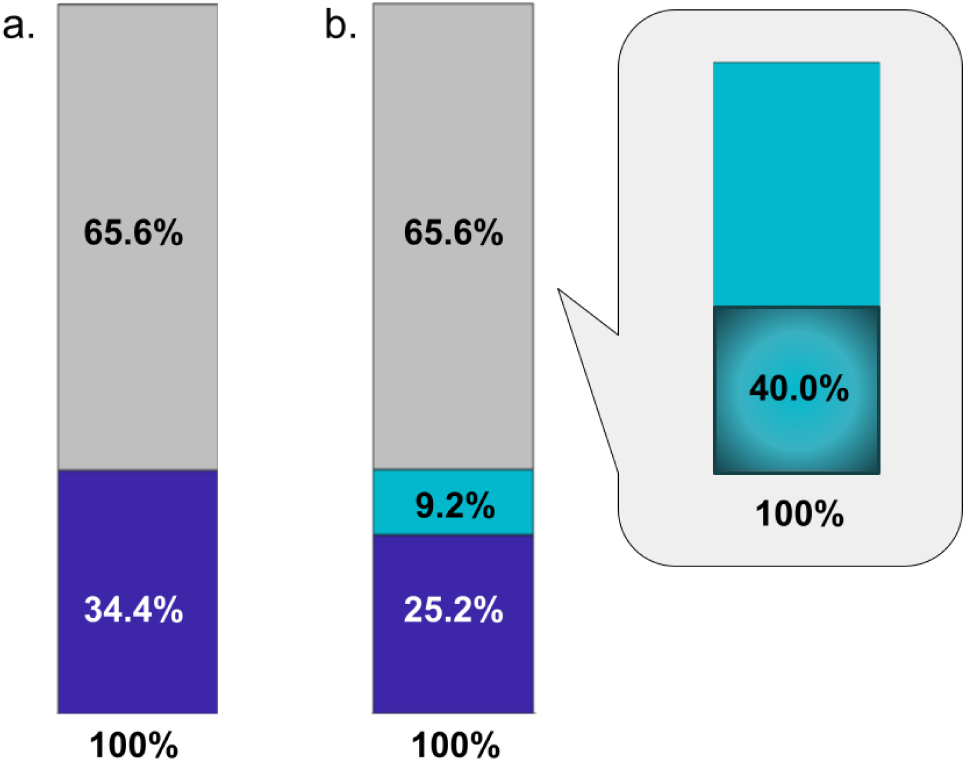
Explained variance of task performance differences. **a)** Total variance (blue) with variance explained by the repeated trials (grey). **b)** The stacked barplot visualizes the variance explained by participant ID only (turquoise), variance explained by the repeated trials (grey), and remaining unexplained variance of the total task performance variance (blue). The second barplot in the grey callout visualized the explained variance by the diameter model (40%) with respect to the total variance explained by individual differences (turquoise).

To better understand the contribution of individual differences to the overall task performance, we first assessed how much of the total variance across all trials could be attributed to participant identity. A simple linear model of all pointing errors with participant identity as a fixed effect resulted in *R*² = .092. Consequently, individual differences explained 9.2% of the total task variance (Figure 9b).

Next, we investigated the link between individual differences in task performance and participants’ self-evaluation of their spatial navigation abilities and spatial strategy preferences. To this end, we modeled the mean pointing error of each participant using participants’ responses to the FRS questionnaire. The 19 questions of the FRS questionnaire are based on a 7-point Likert scale and correspond to three subscales: global-egocentric, survey, and cardinal directions (Münzer & Hölscher, 2011). Therefore, we calculated participants’ mean response to each FRS subscale resulting in the following summary statistics: global-egocentric (*M* = 4.4, *SD* = 0.95, range = 2.5 - 5.8), survey (*M* = 3.85, *SD* = 1.1, range = 1.7 - 6), and cardinal directions (*M* = 2.54, *SD* = 0.65, range = 1 - 6). These mean responses of the three FRS subscales served as the predictors of participants’ mean angular pointing error in a linear model. However, the model resulted in a low R², and none of the predicting FRS subscales were significant (Table 1). Additionally, simple linear regression models of the individual FRS subscales were also non-significant and resulted in very low R² values as well (Table 1). Overall, the FRS questionnaire subscales were not predictive of individual differences in task performance.

**Table 1:** Linear Model: individual differences and FRS questionnaire. This table presents the results of the linear model using the FRS questionnaire scales to predict participants’ mean task performance, i.e., the mean pointing error for each participant (0-180°). In addition to the full model, the results of the simple regression for each predictor of the full model are included.

| **Full model: predicting mean pointing error from all FRS questionnaire scales** |  |  |  |  |
| --- | --- | --- | --- | --- |
| **Fixed effects** | **Estimate** | **SE** | **t** | **p** |
| (Intercept) | 64.405 | 14.097 | 4.569 | .00015 |
| Mean FRS global-egocentric scale | 0.031 | 3.243 | 0.010 | .992 |
| Mean FRS survey scale | -3.008 | 2.847 | -1.057 | .302 |
| Mean FRM cardinal scale | -0.557 | 2.196 | -0.254 | .802 |
| **The model was not significant: $R^2 = .068$, $Adjusted R^2 = -.059$, $F(3,22) = 0.535$, $p = .663$, residual $SD = 14.46$ (DF = 22)** |  |  |  |  |
| **Simple regression results of the individual FRS questionnaire scales** |  |  |  |  |
| **Fixed effect** | **Estimate** | **p** | **$R^2$** |  |
| Mean egocentric/global | -1.658 | .562 | .014 |  |
| Mean survey | -3.097 | .208 | .065 |  |
| Mean cardinal | -0.967 | .656 | .008 |  |

Furthermore, we investigated the link between participants’ visual behavior during spatial exploration and individual differences in task performance. For this purpose, we modeled participants’ mean pointing error with their’ visual behavior during the exploration phase. Specifically, we analyzed participants’ gaze graphs and calculated five global graph measures: number of nodes, number of edges, density, diameter, and graph hierarchy index. These five global gaze graph measures were then used as predictors for participants’ mean pointing error in a linear model. The resulting model was statistically significant with a relatively high *R²* (*R*² = .413, *p* = .044), but out of the five predictors, only diameter was significant, *p* = .010 (Table 2). When testing the variance uniquely explained by each fixed effect, i.e., the reduction of *R*² when removing the component from the model, we also found graph diameter to have the highest uniquely explained variance of .236 (Table 2). As a last step, we modeled each global graph measure as a single fixed effect in a simple linear regression model predicting participants’ mean pointing error. Both graph diameter and density were statistically significant with *p* = .0005 and *p* = .045, respectively (Table 2). However, the density model had an *R*² of .157, while the model using graph diameter to predict the mean performance resulted in a much higher *R*² of .400 (Table 2). Consequently, gaze graph diameter could explain 40% of the inter-participant variance (Figure 9b).

**Table 2:** Linear Model: individual differences and global gaze graph measures. This table presents the results of the linear model using the global gaze graph measures to predict participants’ mean task performance, i.e., the mean pointing error for each participant (0-180°). In addition to the full model, the results of the simple regression for each predictor of the full model are included, in addition to the uniquely explained variance of each predictor in the full model.

| Full model: predicting mean pointing error from global gaze graph measures |  |  |  |  |
| --- | --- | --- | --- | --- |
| Fixed effect | Estimate | SE | t | p |
| (Intercept) | 304.419 | 1439.166 | 0.212 | .835 |
| Number nodes | -1.322 | 6.008 | -0.220 | .828 |
| Number edges | 0.133 | 0.580 | 0.229 | .822 |
| Density | -4031.776 | 16559.996 | -0.243 | .810 |
| Diameter | 9.178 | 3.237 | 2.835 | .010 |
| Hierarchy Index | -0.734 | 5.973 | -0.123 | .903 |
| The model was significant: $R^2 = .414$, $Adjusted\ R^2 = .267$, $F(5,20) = 2.822$, $p = .044$, $residual\ SD = 12.03$ (DF = 20) | | | | |
| Simple regression results of the individual graph theoretical measures and the uniquely explained variance (reduction of $R^2$ of the full model when removing this effect) | | | | |
| Fixed effect | Simple linear regression of individual predictors | | | Unique $R^2$ ($\Delta R^2$) |
| | Estimate | p | $R^2$ | |
| Number nodes | -0.518 | .509 | .018 | 0.0014 |
| Number edges | -0.026 | .054 | .146 | 0.0015 |
| Density | -847.130 | .045 | .157 | 0.0017 |
| Diameter | 10.290 | .0005 | .400 | 0.2356 |
| Hierarchy Index | -7.245 | .232 | .059 | 0.0004 |

## Discussion

In this project, participants answered a spatial strategy questionnaire, explored the virtual city of Westbrook for an extended time while their gaze movements were recorded, and performed an immersive pointing-to-building task. We transformed the recorded data into gaze graphs and applied a graph-theoretical analysis as proposed by previous work (Walter et al., 2022). Supporting the earlier investigation, we reproduce several key properties for the new cohort of participants exploring the new virtual environment: (1) The city of Westbrook is perceived as a whole without easily separable blocks. (2) A few buildings qualify as gaze-graph-defined landmarks that fit typical landmarks in appearance and location. (3) These landmarks are visible during the larger part of the paths taken by the participants in the city. (4) They are preferentially connected with each other, i.e., forming a rich club. Testing the participants’ performance in the pointing-to-building task reveals a large variance. Crucially, the model based on the questionnaire data is not predictive of participants’ task performance. In contrast, global gaze-graph measures, specifically the graph diameter, can explain 40% of the variance attributed to individual differences in the performance of the pointing-to-building task. Thus, gaze graphs appear to be a reliable characterization of visual exploration of virtual cities, and the related graph-theoretical measures allow modeling a substantial fraction of interindividual differences in spatial navigation knowledge.

### Gaze graphs as a representation of visual behavior during free spatial exploration

Recording eye tracking data in virtual reality has many advantages, as it allows for free visual and movement behavior while facilitating the identification of viewed objects; however, the behavioral variability of participants needs to be addressed appropriately in the analysis. Specifically, in this study, participants moved freely within a large area (approximately 75.5 hectares). Consequently, they viewed the city, including all 244 city buildings from various angles, perspectives, and locations throughout the exploration phase. To investigate common patterns in visual attention among participants, we required an analysis approach that could reduce the dimensionality of the recorded eye tracking data to the relevant aspects we were interested in. Since we wanted to understand how the participants learn the spatial layout of the city with respect to the buildings it consists of, we assume that participants learned something about the location and layout of two buildings when they looked directly from one building to the other. Thus, we were interested in participants’ gaze transitions. In general, investigating patterns in gaze transitions has been a focus of eye tracking research for decades (Anderson et al., 2015; Noton & Stark, 1971), and recent publications highlight the importance of sequential analysis of visual attention or scanpaths (Fahimi & Bruce, 2021). However, despite the diverse and varied methods of analysing scanpaths, these methods typically rely on participants being presented the same visual stimuli from the same angle and perspective (Anderson et al., 2015; Duchowski et al., 2010; Fahimi & Bruce, 2021; Goldberg & Helfman, 2010; Raschke et al., 2014). Other gaze-transition analysis approaches like transition matrix and gaze transition entropy focus on the predictability of gaze transitions, or their randomness (Krejtz et al., 2015; Shiferaw et al., 2019). Thus, classical gaze transition analysis approaches are not suited to analyze the gaze transitions between buildings while accounting for large variances in timing, perspective, and visual angle. Therefore, we proposed an alternative graph-theoretical analysis approach in our earlier work (Walter et al., 2022) that transforms the eye tracking data into gaze graphs and subsequently applies a graph-theoretical analysis. In these gaze graphs, each viewed building is represented as a node, and direct gaze transitions between buildings are represented as edges. As a consequence, we represent direct gaze transitions between regions of interest (i.e., buildings) in the graph independent of when or where these transitions occurred in the experiment. Therefore, gaze graphs reflect participants’ path of visual attention during the free spatial exploration while filtering out behavioral variability such as exploration order and differences in visual angle and perspective. Thus, by transforming eye-tracking data into gaze graphs, the highly variable visual behavior resulting from the free exploration becomes comparable, as long as participants explored the same environment. Moreover, the subsequent graph-theoretical analysis can reveal patterns in visual attention that are statistically consistent across participants while also offering insights into the importance and configuration of buildings as reflected by participants’ perception of the city. Consequently, gaze graphs represent relevant aspects of visual behavior dynamics, namely gaze transitions, while filtering out irrelevant behavioral variability, and thus allowing for a comparative analysis of patterns in visual behavior during the free exploration and spatial knowledge acquisition across participants.

To explore whether general characteristics of gaze graphs and visual behavior generalize beyond a single virtual environment, we applied the identical graph-theoretical analysis from our earlier work (Walter et al., 2022) to the data recorded in this experiment within the virtual city of Westbrook. Given the similarities of both experimental designs, the question remained whether these similarities translate to similar gaze graph architectures. A detailed overview of the experimental design and results of this work and our earlier work is listed in Table 3. In addition, for better readability, we will refer to this work with the initials “WB” (for the virtual city of Westbrook) and to our earlier work with the initials “SH” (for the virtual city Seahaven of our earlier work) in the following. In general, there are many similarities between the experimental design of this (WB) study and our earlier SH study. In both studies, participants freely explored a large virtual city with more than 200 buildings for several 30-minute sessions in an immersive VR setting with a head-mounted VR headset and eye tracker. However, the city of Westbrook in this study is larger (75 vs 21.55 hectares, WB and SH, respectively) and contains more buildings (244 vs. 213 buildings, WB and SH, respectively). Also, participants explored the city for 2.5 h vs 1.5 h, WB and SH, respectively. Despite some differences in experimental design, the eye-tracking data of both studies were pre-processed with the same preprocessing pipeline, and the gaze graphs were created with the same logic. Importantly, when comparing the gaze graph architecture in both studies, we found similar patterns in the gaze graph’s structure. Specifically, while the gaze graphs of this study (WB) contained more edges on average compared to the previous study (1295 vs. 883, WB and SH, respectively), the mean percentage of instantiated edges (density) was very similar in both studies (4.56% vs. 4.1%, WB and SH, respectively). Similarly, in the spectral graph partitioning, the average number of edges cut to separate the graph into two components using the sparsest cut approximated by the Fiedler vector differed (122.8 vs. 82.6, WB and SH, respectively), yet the respective mean percentage of the cut edges was very similar (9.39% vs. 9.30%, WB and SH, respectively). Furthermore, we found similar mean Fiedler values (0.353 vs 0.300, WB and SH, respectively), similar distributions of Fiedler vector values, and no modular block structure in the corresponding visualizations. Therefore, we concluded in both studies that the cities were perceived as a whole and could be treated as such for any further graph-theoretical analysis. Summarizing, despite minor differences in exploration duration and city area, we find similar patterns in the gaze graph architecture in both studies, overall highlighting the consistency of gaze graph architecture across both studies.

**Table 3:** Comparison of experimental design and results of the Westbrook (WB) and Seahaven (SH) projects. This table presents the details and results of the current study (Project Westbrook, WB) and our earlier work (Project Seahaven, SH, (Walter et al., 2022)). It includes the details of the experimental design, the pre-processing pipeline, and the results of the graph-theoretical analysis applied in both studies.

|  | Project Westbrook (WB) | Project Seahaven (SH)<br>(Walter et al., 2022) |
| --- | --- | --- |
| **General experiment details** |  |  |
| Number of participants | 26 | 22 |
| Number of buildings in the city | 244 | 213 |
| Exploration time of participants | 150 min | 90 min |
| VR headset | Vive pro eye | Vive |
| Eye tracker | Tobii (Vive pro eye) | Pupil labs VR plugin |
| Mean sampling rate | 90Hz | 30Hz |
| **Pre-processing pipeline** |  |  |
| Properties to determine good data samples / data cleaning | The combined-gaze-validity-bit mask is 3 | The pupil was recognized with at least 50% probability |
| Number of excluded participants due to bad data quality | 0 | 2 |
| Number of remaining participants for data analysis | 26 | 20 |
| Identification of gazes | Continuous cluster of data points on one collider of 266.6ms or longer | Continuous cluster of data points on one collider of 266.6ms or longer |
| Percentage of gazes in the data | 72% gazes on average | 86% gazes on average |
| **Graph partitioning** |  |  |
| Fiedler value (2nd smallest Eigenvalue) | $M = 0.353$, $SD = 0.093$ | $M = 0.300$, $SD = 0.097$ |
| Edges | $M = 1295.1$, $SD = 206.3$ | Mean: 883.5, $SD = 134.2$ |
| Density | $M = 0.0456$, $SD = 0.0066$ | Mean: 0.041, $SD = 0.0058$ |
| % of instantiated edges with respect to all possible edges | $M = 4.56\%$ | Mean: 4.1% |
| % of edges to cut graph into 2 | $M = 9.39\%$, $SD = 1.2\%$ | $M = 9.3\%$, $SD = 1.3\%$ |
| parts |  |  |
| Absolute number of edges to cut graph into 2 parts | $M = 122.8, SD = 29.5$ | $M = 82.6, SD = 19.9$ |
| Approximate edges of each cluster (after partitioning) | Cluster 1:<br>$M = 584.7, SD = 155.7$<br>Cluster 2:<br>$M = 587.7, SD = 212.0$ | Cluster 1:<br>$M = 417.5, SD = 165.8$<br>Cluster 2:<br>$M = 383.35, SD = 137.3$ |
| intra-cluster density (after partitioning) | $M = 0.0873, SD = 0.0133$ | $M = 0.079, SD = 0.012,$ |
| mean inter-cluster density (after partitioning) | $M = 0.0092, SD = 0.0024$ | $M = 0.0083, SD = 0.0020$ |
| **Gaze-graph-defined landmarks** |  |  |
| Node degree of participants averaged over buildings | $M = 10.62, SD = 1.69$ | $M = 8.3, SD = 1.26$ |
| Pairwise inter-participant correlation coefficients | $M = 0.74, SD = 0.06$ | $M = 0.7, SD = 0.12$ |
| Node degree of buildings averaged over participants | $M = 10.62, SD = 4.96$ | $M = 8.3, SD = 3.98$ |
| Hierarchy Index | Ranges between -2.34 and -4.2, | Ranges between -2.44 and -3.77 |
| | $M = -3.1, SD = 0.47$ | $M = -2.91, SD = 0.34$ |
| Node degree threshold to identify gaze-graph-defined landmarks | 20.53 | 16.25 |
| Number of gaze-graph-defined landmarks | 5 | 10 |
| Percentage of buildings in the city identified as gaze-graph-defined landmarks | 2.05% | 4.69% |
| **Walked area and visibility** |  |  |
| Percentage of the walked area where at least 2, exactly 1, no gaze-graph-defined landmarks were visible | 2 or more: 6.62%<br>1: 21.86%<br>0: 71.52% | 2 or more: 39.1%<br>1: 32.7%<br>0: 28.1% |
| Time spent in areas where at least 2, exactly 1, no gaze-graph-defined landmark was visible | 2 or more: 15.14%<br>1: 31.82%<br>0: 53.04% | 2 or more: 53.2%<br>1: 19.4%<br>0: 27.4% |
| **Rich club analysis** |  |  |
| Threshold of max node degree of nodes considered for the adjusted rich club coefficient (mean + std) | 15 | 13 |
| Mean/std frequency of the buildings appearing in the rich club with the max node degree threshold | $M = 6.2, SD = 7.6$ | $M = 3.7, SD = 5.3$ |
| Threshold to identify the rich club buildings (count mean + 2* std) | 21.38 (max count = 26) | 14.26 (max count = 20) |
| Number of rich club buildings | 14 (including all 5 gaze-graph-defined landmarks) | 14 (including all 10 gaze-graph-defined landmarks) |

### Gaze-graph-defined landmarks

Based on the graph-theoretical analysis of eye tracking data, we identified five gaze-graph-defined landmarks in Westbrook following our definition from Walter et al. (2022) and assessed their characteristics. In general, gaze-graph-defined landmarks are buildings that are viewed exceptionally often directly before or after viewing other buildings in the city. In line with this definition, all five gaze-graph-defined landmarks exceeded the two standard-deviation distance from the population mean node degree, while all gaze graphs displayed a clear hierarchical structure with respect to their node degree distribution. Additionally, all of these five buildings had a distinct appearance (e.g., a distinct facade, painted street art, one is a church, and one is a silo) and were relatively free-standing, making them visible from multiple locations. Four of the five gaze-graph-defined landmarks were located next to crossings and overall centrally located in the city, while the fifth gaze-graph-defined landmark was a tall silo at the edge of the city. These characteristics correspond with our findings that gaze-graph-defined landmarks were viewed from a much larger city area compared to most other buildings. Also, at least one of the gaze-graph-defined landmarks was visible in nearly one-third of the traversed city, and participants also spent nearly half of the experiment time exploring these areas. Altogether, the identified gaze-graph-defined landmarks match characteristics of both global and local landmarks (Chan et al., 2012; Lynch, 1960; Steck, 2000; Yesiltepe et al., 2021). Furthermore, we investigated the visibility and interconnectedness of gaze-graph-defined landmarks. In general, high node degree buildings, defined by a node degree of one-sigma-distance from the population mean, formed the strongest interconnections within the gaze graph when compared to matching random graphs, i.e., the rich club. Moreover, when accumulating which buildings appear most often in the rich club, we identified 14 buildings, including all gaze-graph-defined landmarks. Since both measures depend on a population-based node degree threshold, the connection of the gaze-graph-defined landmarks and the buildings appearing most often in the strongest rich club was expected. Nevertheless, our findings showcase gaze-graph-defined landmarks to be preferentially connected with each other and form a rich club across participants. Overall, the gaze-graph-defined landmarks matched characteristics of both global and local landmarks, as well as being preferentially connected with each other and forming a rich club.

To investigate the consistency of gaze-graph-defined landmark characteristics, we compare the identified gaze-graph-defined landmarks with our earlier work (Walter et al., 2022). On the one hand, the gaze-graph-defined landmarks in both cities (SH and WB) display similar characteristics. They all have a distinct appearance, are located next to crossings, and are relatively near the city center. Additionally, we identify the same number of buildings in both studies that appear most often in the rich club, including all gaze-graph-defined landmarks, despite using a different cut-off threshold to calculate the mean-adjusted rich club coefficient. Thus, the gaze-graph-defined landmarks in both studies were preferentially connected with each other and formed a rich club. On the other hand, we identified more gaze-graph-defined landmarks in Seahaven than we did in Westbrook (10 vs 5, SH and WB, respectively), despite Seahaven being the smaller city with fewer buildings. Similarly, we found some differences in the size of the traversed area in which the gaze-graph-defined landmarks were visible. Specifically, the traversed area in which at least one gaze-graph-defined landmark was visible was smaller in the current study compared to our earlier results (71.8% vs 28.5%, SH and WB, respectively). Correspondingly, we found a smaller traversed area in which at least two or more gaze-graph-defined landmarks were visible simultaneously (39.1% vs 6.6%, SH and WB, respectively). This pattern is mirrored in the fraction of experiment time participants spent in these city area types. Specifically, participants spent less time in city areas where at least one gaze-graph-defined landmark was visible (72.6% vs 46.9%, SH and WB respectively), and we found an even larger difference when comparing the fraction of experiment time participants spent in city areas where two or more gaze-graph-defined landmarks were visible simultaneously (53.2% vs 15.1%, SH and WB respectively). While these differences might be related to Westbrook only having half as many gaze-graph-defined landmarks compared to Seahaven, while simultaneously being the larger city with more buildings, it is also worth noting that the calculation in Westbrook used the raw viewing data, while the analysis in Seahaven applied a kernel convolution, resulting in a morphological dilation. Nevertheless, despite the larger differences in size of the traversed area types, participants still spent nearly half of the experiment time in areas where at least one gaze-graph-defined landmark was visible. Overall, gaze-graph-defined landmarks were consistent in their characteristics and properties across different virtual cities and experimental conditions.

Interestingly, while gaze-graph-defined landmarks match characteristics of landmarks as described in the literature, they were not originally intended as landmarks when designing and building the city of Westbrook. Originally, the city of Westbrook was designed with four tall and distinct-looking buildings that were placed in all four corners of the city and intended to serve as orientation markers and landmarks for participants (Schmidt, König, Dilawar, Sánchez Pacheco, et al., 2023). However, only one of these intended landmarks was also identified as a gaze-graph-defined landmark in our analysis: the high-silo building. This building was the only gaze-graph-defined landmark that was intended as a landmark in the original city design and was also the only gaze-graph-defined landmark that was located at the edge of the city, whereas all other gaze-graph-defined landmarks were located centrally within the city. The remaining three intended landmarks had much lower mean node degree centrality values compared to the gaze-graph-defined landmarks, and therefore, they did not serve as important nodes in the gaze graphs. While further research and more analysis will be necessary to investigate the role of these intended landmarks, a potential explanation could be related to the location of gaze-graph-defined landmarks. In this study, as well as our earlier work (Walter et al., 2022), we find gaze-graph-defined landmarks to be centrally located in a city next to crossings, whereas the intended landmarks were placed at the corners and edges of the city, which could affect their visibility as well as their usefulness as orientation markers, since they were not located directly at decision points. In general, these findings highlight the specific nature of the definition of gaze-graph-defined landmarks. While the designed landmarks were assigned their intended function as a landmark, which ultimately did not translate into behavior, gaze-graph-defined landmarks are defined based specifically on general patterns in visual behavior across all participants and therefore are a data-driven definition. Overall, the identification of gaze-graph-defined landmarks is based on a data-driven definition and thus rooted in the participants’ visual perception of the city. Simultaneously, gaze-graph-defined landmarks match the characteristics of typical local and global landmarks as suggested by the literature and consequently highlight the utility of using such data-driven definitions in spatial navigation research.

Despite the ubiquitous presence of landmarks in spatial navigation research, the role and function of landmarks is still highly discussed. In the case of gaze-graph-defined landmarks, their characteristics suggest that they could serve several functions during spatial exploration and navigation, such as their appearance, placements in the city, visibility, interconnectedness, and others that were suggested by earlier literature. For example, not only were the majority of gaze-graph-defined landmarks located next to complex crossings, they were also visible in more of the walkable area of the city compared to most other buildings, thus they would be suited to serve as orientation markers at decision points (Chan et al., 2012; Mallot & Gillner, 2000; Richter & Winter, 2014; Wang et al., 2014). Moreover, in 65% of the traversed area, at least one gaze-graph-defined landmark was visible, offering the basis of directional cues and beacon navigation (Chan et al., 2012; Ekstrom & Isham, 2017; Richter & Winter, 2014; Waller & Lippa, 2007). Furthermore, in 25% of the traversed area, two or more gaze-graph-defined landmarks were visible; thus, these areas would allow participants to triangulate their position (Ekstrom & Isham, 2017). Given these characteristics, participants could have potentially utilized gaze-graph-defined landmarks as associative cues at decision points, as an orientation or a directional cue, as beacons, as reference or anchor points, or for guidance or triangulation (Chan et al., 2012; Ekstrom et al., 2018; Ekstrom & Isham, 2017; Mallot & Gillner, 2000; Parra-Barrero et al., 2023; Richter & Winter, 2014; Waller & Lippa, 2007; Wang et al., 2014; Yesiltepe et al., 2021). Moreover, since the gaze-graph-defined landmarks were also preferentially connected with each other, forming a rich club, they might not only serve as important navigation markers but possibly also anchor participants’ mental map or allocentric representation, i.e., survey knowledge. In that sense, the interconnected “rich club” of gaze-graph-defined landmarks could serve as a protomap, and thus, represent an important step in spatial knowledge development and the formation of a mental map. However, without investigating the visual behavior during the spatial exploration and the development of the gaze graph over the course of the spatial exploration in more detail, these theories will remain speculative. Nevertheless, the identification of gaze-graph-defined landmarks in two different virtual environments and experimental setups that are consistent in their properties supports the argument that the data-driven definition of gaze-graph-defined landmarks captures an important aspect of visual behavior related to landmarks. Consequently, further research focusing on gaze-graph-defined landmarks could offer new insights into their role in spatial navigation and their function in the development of spatial knowledge and the mental map. Altogether, gaze-graph-defined landmarks show consistent properties that are in line with multiple functions in spatial navigation, as suggested by the literature. As a consequence, gaze-graph-defined landmarks appear to play an important role in spatial exploration, navigation, and the formation of spatial knowledge and the mental map.

### Individual differences in spatial navigation performance

The immersive pointing-to-building task participants performed after the free exploration of the virtual city revealed considerable individual differences in mean task performance, and thus, in spatial knowledge and spatial navigation ability. Specifically, the pointing-to-building task assesses participants’ survey knowledge (Schmidt, König, Dilawar, Sánchez Pacheco, et al., 2023) and mental map accuracy (He et al., 2023; Hegarty et al., 2023). In general, individual differences in spatial navigation tasks are a well documented phenomenon in spatial navigation research (Boone et al., 2019; Ekstrom & Isham, 2017; Gramann, 2013; He et al., 2023; Hegarty et al., 2018, 2023, 2023; Ishikawa, 2023; Ishikawa & Montello, 2006; Maxim & Brown, 2023; Newcombe, 2018; Newcombe et al., 2023; Schinazi et al., 2023; Spiers et al., 2023; Weisberg et al., 2014; Weisberg & Newcombe, 2014, 2016, 2018; Wolbers & Hegarty, 2010; Zanchi et al., 2022). Since our experimental design directly combined the free exploration, i.e., spatial knowledge formation, with a task to assess participants’ spatial knowledge, it allowed us to directly link participants’ visual behavior during spatial knowledge acquisition with their navigation performance. Modeling participants’ mean performance with several global gaze graph measures resulted in a relatively high R², thus, global patterns in visual behavior could explain 41% of the variance between participants’ mean performance. However, only the gaze graph diameter was a significant predictor and had the highest uniquely explained variance. Moreover, a simple linear regression of gaze graph diameter and mean pointing error resulted in a comparable *R²*, indicating that diameter alone explains most of the variance captured by the full model. While a simple linear regression with gaze graph density was also significant in predicting participants’ mean task performance, its R² was much smaller compared to gaze graph diameter, and its uniquely explained variance in the full model was also small. Consequently, gaze graph diameter not only contributes the highest uniquely explained variance, but also appears to capture shared variance with other metrics, specifically gaze graph density, reducing their additional explanatory value. Overall, global patterns in visual behavior during spatial learning and exploration explained 41% of individual differences in spatial navigation performance. In contrast, using self-reported spatial navigation abilities and strategy preferences from the FRS questionnaire did not significantly predict participants’ mean task performance. Since self-report questionnaires have a long-standing tradition in human-based spatial navigation and the FRS has not only been validated with real-world spatial tasks (Münzer & Hölscher, 2011), but has also been predictive in other studies, e.g. (Brunyé et al., 2018; Löwen et al., 2019; Münzer & Stahl, 2011; Münzer & Zadeh, 2016), the question arises, why the FRS questionnaire is not able to model the task performance in this study. While other research also reports weak or no significant correlations, in line with our findings (Credé et al., 2019; König et al., 2019, 2021; Wunderlich & Gramann, 2021), the referenced research also varies widely in tasks and experimental conditions, which is a common issue in spatial navigation research investigating individual differences (Newcombe et al., 2023) and could explain the non-significance of the self-assessment questionnaire here. In addition, to our knowledge, this is the first study that combines a long free exploration phase of a large urban city in immersive VR with an immersive testing scenario in the same virtual environment. Thus, it is possible that the FRS questionnaire does not generalize to the more naturalistic conditions of this study. Interestingly, Schmidt et al. (2023) did find an increase in the FRS scores after the experimental group trained with the feelSpace belt compared to their initial answers and compared to the control group. These findings suggest that while the FRS appears to be sensitive to training-induced shifts in self-assessed spatial abilities and strategy use, it was not able to predict the pointing-to-building performance in our sample and did not account for between-participant differences. Consequently, global patterns in visual attention during spatial exploration and knowledge acquisition predict individual differences in spatial task performance, whereas the self-report FRS questionnaire does not.

While we can explain a substantial portion of the between-participants mean performance differences via participants’ visual behavior during spatial exploration of the city, these findings should be interpreted in the context of other systematic patterns in task performance. Specifically, we found that participants’ task performance significantly increased during the repeated trials of the task, averaging a 5.1° decrease in the pointing error. While this improvement might be related to incidental exposure to new visual information during the task, it could also just reflect general learning effects and task-related adaptation. Without further assessment, we have to assume that multiple factors contributed to participants’ performance improvement and not only newly acquired spatial knowledge. Therefore, we included all task trials (both initial trials and repetitions) in the further performance analysis and modeling approach. In addition, we assessed the intra-participant variance between each unique start-target-building combination to estimate the size of the variance attributed to participants’ inconsistencies in task performance. We found that nearly two-thirds of the total variance can be attributed to participants’ performance differences between the initial and repeated trials, leaving about one third of the variance as an upper bound for how much variance can be captured by models based on participants’ stable abilities or exploration behavior. Furthermore, while participant identity only explained about 9% of the total variance, this small portion of the total variance has to be interpreted with respect to the large within-participant variability that accounts for the majority of the total variance. Therefore, our findings indicate that slight performance improvement during the repetitions and substantial within-participant fluctuations coexist with stable between-participant differences. Nevertheless, more research will be required to investigate the substantial within-participant fluctuations and underlying mechanisms related to participants’ improved performance during the repetitions. Overall, despite the small performance increase during the repeated trials and large within-participant fluctuations, we found reliable between-participant differences, which can be explained by global gaze graph measures, thus offering a robust marker of individual differences in spatial navigation performance.

Moreover, our results offer a new perspective into the underlying factors contributing to individual differences in spatial navigation. In general, multiple factors have been associated with individual differences in spatial navigation, including the environment of the area a person grew up in (Coutrot et al., 2022), different languages and cultural differences (Coutrot et al., 2018; Haun et al., 2011; Hund et al., 2012; Spiers et al., 2023), and cognitive abilities (Weisberg & Newcombe, 2016). Overall, the embodiment in the environment appears to shape spatial navigation preferences, strategies, and abilities (Gramann, 2013), although also stress and anxiety (Maxim & Brown, 2023; Varshney et al., 2024), motivation (Schinazi et al., 2023) and instructions (Boone et al., 2019) appear to modulate individual differences in spatial navigation performance. Importantly, our results indicate that the underlying processes contributing to the observed individual differences in this study were reflected in participants’ visual behavior during spatial knowledge formation. Thus, studying visual behavior during exploration and tasks in more detail offers new insights into the underlying processes and behavioral patterns that contribute to the observed individual differences. Overall, our findings support that studying vision offers access to new insights into individual differences in survey knowledge and their allocentric representation or mental map, therefore offering new opportunities to investigate the factors contributing to individual differences during spatial knowledge acquisition.

### Vision in spatial navigation

In general, our findings in this study offer new insights into the role of vision during spatial exploration and individual differences in navigation performance, consequently highlighting the role of vision for spatial navigation research. Importantly, these findings are in line with a long history of publications highlighting the importance of vision for spatial navigation (Chan et al., 2012; Ekstrom, 2015; Ekstrom & Isham, 2017; Foo et al., 2005). Moreover, combining an immersive and free exploration phase of an environment with an immersive pointing-to-building task offers a unique opportunity to link visual behavior during spatial knowledge acquisition directly with navigation performance. Especially, considering that the self-report questionnaire was not predictive of individual differences, whereas global patterns in visual behavior during spatial exploration were able to explain 41% of interindividual variance. Nevertheless, a considerable portion of the total variance in task performance remains unaddressed, and further analysis will be required to investigate which factors contribute to the observed performance differences. Moreover, while our results highlight the benefit of transforming eye tracking data into gaze graphs and, thus, condensing the complex and high-dimensional eye tracking data into one graph for each participant, this process also eliminates more fine-grained information that could be crucial to understand more specific patterns in visual behavior during spatial exploration, learning, and knowledge formation. Furthermore, so far, we have exclusively focused on the visual behavior of participants as one visual network combining all 2.5 hours of visual exploration in one gaze graph. In the future, investigating visual behavior over the course of the full exploration and combining vision and movement analysis could enable a more complete assessment of behavioral patterns during spatial exploration. For example, two recent publications investigated movement and walking behavior at decision points during spatial exploration and linking explorative patterns with the space syntax of the environment (Brunec et al., 2023) and human agents (Sánchez Pacheco et al., 2025). Nevertheless, the role of vision in these processes remains to be determined. Therefore, investigating the development of graph-theoretical measures, gaze-graph-defined landmarks, and other features of visual attention over the full exploration time in combination with participants’ movement patterns presents a promising approach to create a better understanding of spatial learning, knowledge formation, and landmark functionality. In addition, examining participants’ visual behavior during the pointing task, especially during the reorientation after being teleported to a new location, with respect to gaze-graph-defined landmarks and other orientation markers, presents a promising opportunity to investigate active orientation processes, specifically with a focus on individual spatial knowledge differences. Since these future research directions investigations are more time-sensitive with respect to gaze transitions compared to the analysis applied in this work, the definition of gazes might need to be adapted, and a more fine-grained detection algorithm should be applied in the pre-processing, compared to the fixed threshold of cluster duration applied in this work. A possible velocity-based gaze detection algorithm adaptation for eye tracking data in virtual reality that would fit these requirements was proposed by Nolte et al. (2024). Overall, this study demonstrates the importance of vision and outlines how applying a complex and naturalistic experimental design with freedom of movement and eye tracking recording offers new insights into spatial knowledge acquisition and visual behavior with regard to individual differences. Moreover, given that the self-report questionnaire was not predictive of participants’ individual performance differences, this study also highlights the importance of using more objective and observational methods to investigate spatial navigation. Altogether, our findings demonstrate that gaze graphs and the graph-theoretical analysis of eye tracking data are an informative analysis of visual behavior in spatial exploration and navigation that highlights the importance of vision for spatial navigation research and outlines the potential future research holds in applying and expanding the presented analysis approach.

To conclude, visual behavior during the free exploration of the virtual city explains individual differences in navigation performance, and this study demonstrates the consistency and potential of the graph-theoretical analysis approach of eye tracking data. Specifically, gaze graphs are a useful representation of the complex visual behavior with freedom of movement and allow to investigate similarities and differences across participants. The characteristics of gaze-graph-defined landmarks are in line with multiple functions in spatial navigation, as suggested by the literature, and appear to play an important role in spatial exploration, the formation of spatial knowledge, and the mental map. In addition, the consistency of results and properties of gaze graphs and gaze-graph-defined landmarks also support the validity of the graph-theoretical analysis. Moreover, the global gaze-graph measures could explain a large proportion of the individual differences in spatial task performance. Therefore, we conclude that differences in visual behavior during spatial knowledge acquisition explain individual differences in spatial navigation performance. Consequently, our results underline the importance of vision and eye tracking methodology for spatial navigation research. While some questions remain unaddressed, future research applying and expanding the graph-theoretical method promises new insights into spatial knowledge formation, landmark functionality, orientation processes, and spatial navigation. Overall, this study showcases the important role of vision for spatial navigation research, offers a new perspective on individual differences in spatial navigation and potential landmark functionality, and demonstrates the potential of the graph-theoretical analysis approach for investigating vision in a naturalistic and immersive spatial navigation paradigm with freedom of movement.

## Methods

### Ethics statement

The study was conducted in accordance with the Declaration of Helsinki and approved by the Ethics Committee of the University of Osnabrück (protocol code: 4/71043.5, date of approval: 5 January 2021). Informed consent was obtained from all subjects involved in the study.

### Data availability

The anonymized eye and motion tracking dataset analyzed in this study can be downloaded here: https://osf.io/qcn67 (Schmidt, Walter, et. al., 2025). The project documentation, the VR exploration and VR task assessment builds for replication or incorporation in future VR experiments, and the performance data and questionnaire responses can be downloaded here: https://osf.io/32sqe (Schmidt, König, Dilawar, Pacheco, et al., 2023). All pre-processing and analysis scripts are available at the project’s GitHub repository https://github.com/JasminLWalter/gaze-graphs-in-spatial-navigation, which is also archived with Zenodo and assigned a permanent identifier (Walter, 2025).

### Study design and participants

The work presented here is part of a larger study that followed a mixed-methods design, aiming to assess behavioral and qualitative self-reported changes related to spatial knowledge acquisition in a virtual environment. The behavioral results of this study, including a comparison to a group of participants that were trained to use an augmented exploration, are published in Schmidt et al. (2023). As the present study focuses on general behavioral patterns during spatial exploration and navigation, only the data from the control group were considered in the current analysis. Participant exclusion criteria involved age below 18 or above 39, past occurrence of sea or motion sickness, past or current substance abuse, use of psychotropic drugs, medical abnormalities that could impact cognitive functions, and history of neurological conditions or head injury. After consideration of all these factors, a total of 28 participants were recruited for the control group. Two participants dropped out during the first session due to acute motion sickness. The final sample considered for analysis, therefore, contained 26 participants, aged between 18 and 35. As an incentive for participation, all individuals received a monetary reward or university-internal participation credits, according to their preference.

### Virtual reality setup and data collection

The experiment was conducted in an immersive virtual reality using the Vive Pro Eye headset (Resolution: 1440 x 1600 pixels per eye (2880 x 1600 pixels combined), Refresh rate: 90 Hz, Field of view: 110 degrees) and Valve Index controller (Figure 1a). Participants were located on a swivel chair that allowed them to fully rotate their bodies (Figure 1a). During the exploration phase of the experiment, participants were able to walk in the environment using their controller. Unlike typical virtual reality setups, the direction of walking, i.e., forward direction, was determined by the HTC tracker placed at the center of the participant’s chest instead of the headset orientation. Therefore, participants were free to rotate their head without any interference with their walking direction, providing more naturalistic conditions for the head movement behavior and further minimizing the risk of motion sickness. Participants’ height in the virtual city was set to 1.50 m using the SteamVR room setup, to account for the seated position of the participants and ensure an equal viewing perspective on the objects in the environment.

Eye tracking was performed using the Tobii eye tracker integrated in the Vive Pro Eye headset (accuracy: 0.5°–1.1°, calibration: 5-point, trackable field of view: 110°). The eye tracker was integrated with the Unity VR experiment using the HTC SRanipal interface. The collection of eye tracking data, head and body tracking, controller movements, and the raycast to determine viewed objects was implemented using a coroutine in Unity. However, a programming bug discovered only after data collection affected the waiting time of the coroutine, so that the data collection was performed with a varying sampling rate averaging to 90 Hz. Before every 30-minute exploration session, the full calibration and validation procedure was conducted to ensure eye tracking accuracy. Specifically, the integrated Tobii eye tracker was calibrated using the SRanipal system calibration and validated using a custom-built validation implementation to calculate the validation error. In case the validation error was not below 1° visual angle, the system calibration was repeated until a validation error of 1° or below was achieved. Additionally, throughout the experiment (exploration phase and testing phase), a 1-point validation of the eye tracker was performed every 10 minutes. If the validation error exceeded the threshold of 1° of visual angle, a full system calibration and custom-built validation were performed until the validation error was again below 1° of visual angle.

### Experimental structure

The experiment was structured into five parts:

1. Completing the FRS questionnaire on spatial strategies and preferences
2. Movement training in virtual reality (training environment)
3. Free exploration of the virtual city of Westbrook for 2.5 hours/150 min (separated into five 30-minute exploration sessions)
4. Task training in virtual reality (training environment)
5. Performing spatial navigation tasks in the virtual city of Westbrook

At the start of the experiment, participants were asked to answer the FRS questionnaire on spatial strategies (Münzer & Hölscher, 2011). The FRS questionnaire is a self-tool that aims to measure a respondent’s likelihood to apply certain spatial strategies and their spatial abilities. These are divided into the categories: global-egocentric spatial knowledge, survey knowledge, or cardinal directions. For example, the FRS questionnaire contains questions like “I always imagine the surroundings as a ‘mental map’” or “My sense of orientation is very good.”

Next, participants entered a training environment in virtual reality to learn how to move in virtual reality using their full body rotation and the controller to walk based on their body rotation. They also learned how to take a picture of the environment using their controller. The training was completed once the participants demonstrated fluent walking behavior in VR and took at least one picture.

In the exploration phase of the experiments, participants were asked to explore the virtual city of Westbrook freely for a total of five 30-minute sessions. The exploration sessions took place on 5 separate occasions, but always with at least 6 hours and a maximum of 3 working days in between the appointments. Thereby, we could ensure that all participants received similar learning and consolidation intervals. At the beginning of each 30-minute exploration session, participants were teleported to a central starting location in the virtual city of Westbrook. They could then freely walk around the city’s road network and walkable areas. In order to provide additional motivation to explore the various streets and paths in Westbrook, participants received the task to take pictures of all street art they could find in the virtual environment. The 30-minute exploration sessions were automatically paused every 10 minutes to perform the one-point validation of the eye tracker as described above. Additionally, participants were allowed to request a break at any point they desired during the experiment. In the case of an additional break or if the headset was moved, a full calibration and validation procedure for the eye tracker was performed.

After completing the 2.5 hours of free exploration in the city, participants performed three types of spatial navigation tasks: a task where they were asked to point to a specific building (pointing-to-building task), a task where they were asked to point to cardinal north (pointing-to-north task), and a task where a building was removed from the environment and participants controlled a semi-transparent version of the building with their controller and were asked to place the building at the correct location in the city with the correct orientation. Before performing the tasks in the virtual city, participants entered a training environment, in which they were introduced to the task instructions and performed some test trials to get familiar with the required buttons of the controller to answer the task trials. After successfully answering several training questions of each task type, a full calibration and validation procedure was performed, and participants were teleported to their first task location in the city of Westbrook.

The work reported in this publication is based on the performance data of the pointing-to-building task only. However, it should be noted that while the pointing tasks were performed first, the order of the pointing-to-building and pointing-to-north tasks was randomized. Consequently, some participants performed a pointing-to-north task before the pointing-to-building task (Schmidt, König, Dilawar, Sánchez Pacheco, et al., 2023).

The pointing-to-building task is aimed at measuring participants’ spatial knowledge, specifically their survey knowledge (Schmidt, König, Dilawar, Sánchez Pacheco, et al., 2023) and mental map accuracy (He et al., 2023; Hegarty et al., 2023). To maintain the naturalistic conditions and immersiveness of the exploration phase, participants performed the pointing task within the same environment by being teleported to different locations in the city and asked to point to the task buildings (Figure 7a). The task consisted of eight task buildings that all featured street art but varied in size and appearance (Figure 7c), and eight corresponding task locations that were directly located in front of a task building. The task buildings and locations were evenly distributed over the city area (Figure 7b), and at each task location, none of the other seven task buildings were visible. During the task, participants were teleported to the eight task locations in the city, placed directly in front of the “start” building of the trial, and asked to point to the remaining seven task buildings, the pointing targets. The order of task locations and pointing “target” buildings was randomized. After completing all seven trials in one location, participants were teleported to the next random task location. Therefore, each participant performed 56 unique “start-target-building” pointing trials. After completing all 56 pointing trials, the participants repeated all trials again in a newly randomized order to assess participants’ consistency in the pointing task. Therefore, participants performed a total of 112 pointing-to-building trials. During the task, all participants were set to the same physical height, and translational movement was disabled, whereas rotational movements remained enabled to ensure that the available visual information was consistent across participants. In addition, participants were always rotated to a random degree during the teleportation process in the virtual environment. At each task location, the “target” building of each trial was displayed in a small square in the top center of the HMD, which could be enlarged with a button press. To point towards the “target” building, participants lifted their right or left controller into the desired direction (Figure 7a). The pointing direction was displayed in the virtual environment in real time in the form of a green beam. By pressing the trigger button of the controller, participants were able to freeze the pointing beam. When they were satisfied with the pointing direction, they could confirm their response by pressing the trigger button again and continuing to the next trial. Alternatively, they could correct the pointing direction by unfreezing the beam using the “B” button on the controller. After participants confirmed their pointing response to a trial, the next “target” building would be displayed, or they would be teleported to the next task location.

### The virtual town of Westbrook

The virtual town of Westbrook has a size of 75.5 hectares (996 m x 758 m) and features 232 unique residential or commercial building types, a castle, a farm, a school and a silo, as well as several garages (5 were counted as separate buildings as they were free-standing, all others were regarded as part of the respective building), and two cranes (Figure 1b-1c). This resulted in a total of 244 unique buildings and objects of interest. The environment additionally features various parks and construction sites with typical features such as benches, trees, fences, barriers, and scaffolding. Next to a bus stop, the town of Westbrook also features a basketball court and a realistic road network consisting of streets and paths in various sizes, inspired by the city of Baulmes in Switzerland (though heavily modified). In order to limit the amount of movement possibilities while preserving free exploration of the virtual town, the movement of participants was limited to the road network and shortcuts over grass and small trails, as visible in participants’ walking paths (Figure 5a). To this end, we equipped all roads and even smaller paths with curbs, steps, or fences as a natural visual barrier indicating movement limits. To minimize potential social influences on behavior, no virtual non-player characters were included in the virtual town. However, to enhance ecological realism, we included static bicycles and cars. To avoid people using the clouds or the sun as orientation, we modified a cloudy but relatively bright skybox to include the same mirrored image of clouds three times and disabled shadows from being rendered. The aim of the virtual town of Westbrook is to provide an environment with optimal conditions to obtain data about spatial knowledge acquisition in a controlled setting that still feels naturalistic to the research participant.

### Data pre-processing and graph-theoretical analysis

The eye tracking and head tracking information were already synchronized and combined during the experiment runtime to calculate a gaze vector in viewing direction. In addition, a raycast was performed using the gaze vector in viewing direction, and all colliders that were hit by the ray were calculated. The hit colliders were then sorted according to their distance from the gaze origin to the hit point on the collider, and the two colliders closest to the participant were selected and saved. Importantly, all eye tracking, unity-specific, and raycast variables were synchronized and saved during runtime.

As a first step in the pre-processing pipeline, the first hit collider was substituted with the second hit collider in specific cases where the following predefined conditions were met. In case the first hit collider was transparent (e.g., invisible barriers for experiment functionality) or see-through (e.g., wire-mesh fence), the first hit collider was replaced with the second hit collider. Due to the amount of natural objects in the virtual city that were often surrounded by colliders that did not exactly match the corresponding visible object (trees, bushes, fences etc.), the first collider was also replaced with the second hit collider, in case the first hit collider belonged to a natural object and the second hit collider correspond to a building in the city. An example of such a case would be a data point in which the first hit collider corresponded to the scaffolding around a building and the second hit collider corresponded to the building itself. Next, the eye tracking data was cleaned based on the SRanipal variable “combined validity bitmask”, which is a system evaluation measure of the eye tracking quality. Data samples that did not have a combined validity bitmask value of 3 were marked as invalid. In invalid data rows, all data variables based on the eye tracking or raycasting information were marked as NaN and thus removed from the analysis. On average, 4.75% of the data was removed during this cleaning process.

The subsequent pre-processing, i.e., interpolation and gaze identification, was adapted from our earlier work (Walter et al., 2022). Specifically, consecutive data samples with the same hit collider were combined into data clusters. An interpolation was performed on the cleaned data samples only if the data cluster of cleaned data, i.e., a cluster of NaN values created during the cleaning process, had a duration of 266.6 ms or shorter and was located between data clusters of the same collider. During the interpolation, the missing data cluster collider information was interpolated and combined with the surrounding data clusters. An example of this principle would be the following: the eye tracking data contains a hit collider cluster on building 01, followed by a cluster of NaN-values (smaller than 266 ms), followed by another cluster on building 01, resulting in a combination of all 3 data clusters into one data cluster on building 01 during the interpolation process. Following the interpolation process, valid gazes were identified in the data. Specifically, data clusters were classified as gazes if the data cluster of a collider had a duration of at least 266.6 ms. Subsequently, these gazes serve as the basis for the creation of the gaze graphs. On average, 72% of the data was classified as gazes.

Finally, the gaze graphs were created for each participant, based on the identified gazes in the pre-processing, again following the creation process and definitions of our earlier work (Walter et al., 2022). In the gaze graph, nodes represent buildings viewed by the participant. Therefore, if the data of a participant included at least one gaze on a building, a node representing this building was created in the corresponding gaze graph. An edge between two nodes was created when a participant made a direct gaze transition between the two buildings corresponding to the respective graph nodes. Therefore, edges in the gaze graph represent the direct gaze transitions between two viewed buildings. Note that gazes on non-building objects (e.g., road, trees, cars, etc.) were ignored in this process. Therefore, if a participant directed their gaze from a building to a tree, followed by a gaze at another building, the graph would contain an edge between these two buildings only. Importantly, the edges are undirected and unweighted. Thus, if a specific edge already exists in the gaze graph, it is not created a second time. Furthermore, in case the continuous eye tracking data was interrupted, for example, by a remaining missing data cluster (i.g, data loss that was too large to be interpolated) or a temporary break (interruptions of the spatial exploration), the gaze graph was disconnected as well, and no edge was created in the gaze graph. Such temporal breaks included the validation of the eye tracker every 10 min, the longer breaks after every 30 min recording, and additional interruptions or breaks that occurred during the recordings.

Please note that the created gaze graphs are undirected. That is, the directionality of the gaze transitions, i.e., whether the participant gazed from one house to another or the other way around, is not represented in the graph. Also, since the gaze graphs are not weighted, the number of times a specific gaze transition between buildings occurred is also not represented in the gaze graphs. A detailed description of the gaze graph creation process with additional visualizations and mock-up graphs with detailed descriptions of more complex graph-theoretical measures can be found in (Walter et al., 2022).

## Acknowledgments

We gratefully thank Philip Spaniol, Nora Maleki, and Linus Tiemann for their technical support as research assistants in the development of the VR environment Westbrück. As a team, they provided outstanding contributions to the technical development of a realistic urban VR environment and the corresponding VR tasks, which were customized for the needs of spatial navigation research. Similarly, we want to thank the students Mohammad Abdul Munem and Melissa-Andrea Sarria-Mosquera for their support with the execution of exploration and task assessment sessions. In addition, we gratefully acknowledge the support by the Research Training Group Computational Cognition, which is funded by the Deutsche Forschungsgemeinschaft (DFG, German Research Foundation) – GRK2340. In addition, this research was funded by the EU Horizon 2020 (MSCDA) research and innovation program under grant agreement No. 861166 (INTUITIVE). The funders had no role in the study design, data collection and analysis, decision to publish, or preparation of the manuscript.

